# Operating regimes in a single enzymatic cascade at ensemble-level

**DOI:** 10.1101/564922

**Authors:** Akshay Parundekar, Girija Kalantre, Akshada Khadpekar, Ganesh Viswanathan

## Abstract

Operating regime of a single enzymatic cascade such as ubiquitously conserved MAPK building block provides insights into the nature of the sensitivity of the steady-state dose response that maps the upstream kinase and the downstream activated substrate concentrations. Steady-state response can be viewed either at a single-cell level in an ensemble of cells or at population-average level. Four operating regimes, viz., hyperbolic, threshold-hyperbolic, signal transducing and saturated, have been identified using the population-average level dose response curve. However, cell-to-cell variability exists in enzymatic cascades. This variability is captured using reporter based experimentation at ensemble level which permits detection of snapshot(s) of ensemble-level distribution of phosphorylated proteins. As a result, often the corresponding underlying steady-state dose response curve may not be available. We consider the question if the underyling operating regime can be directly inferred from ensemble level snapshot upstream kinase (input) and downstream phosphorylated substrates (response) distributions. In order to address this question, we use mathematical model of a single enzymatic cascade based on quasi-steady state approximation superimposed with an input distribution constrained by single-cell level experimental measurements of the MAPK cascade in Jurkat E6.1 cells stimulated with Phorbol Myristate Acetate (PMA). We prove that, under steady-state conditions, a monotonic relationship between the R_IQR_ (=ratio of the inter-quartile range of the response and input distributions) and R_m_ (=ratio of medians of the two distributions), both of which are experimental observables, can be used to identify the underlying operating regime. We also show that the identification of the unimodal vs bimodal nature of the response distributions can further lead to identification of the potential parameter range in the planes of Michaelis-Menten constants *K*_*1*_ and *K*_*2*_, the two key parameters that dictate the operating regimes. We implement the proposed method on the stimulus strength dependent steady-state single-cell level pMEK (input) and pERK (response) distributions in Jurkat E6.1 cells treated with PMA. While cells stimulated using low concentrations of PMA are likely to operate in hyperbolic regime, those exposed to higher concentrations may lie in signal-transducing regime.

**Author’s summary:** Single enzymatic cascade, a ubiquitously found key building block in biological signaling networks, exhibits different steady-state behaviour at population (ensemble) level compared to population-averaged response. Detection at ensemble level typically achieved by non-plasmid based reporter permits snapshot fluorescence distribution measurement. We ask if there are signatures of fluorescence distribution of input kinase and response activated protein that can help decipher the underlying dose-response belonging to four distinct operating regimes a single enzymatic cascade exhibits. Based on simultaneously measured snapshot input (pMEK) and response (pERK) protein levels in an ensemble of PMA stimulated immortalized cancer (Jurkat E6.1) cells, we superimpose pMEK data-guided upstream kinase distribution capturing cell-to-cell variability on the steady-state Michaelis-Menten (MM) kinetic model. Following extensive MM constants sampling and simulations, we systematically identify that monotonicity between R_IQR_, the ratio of the inter-quartile range of response and input distributions, and R_M_, the median ratio of the two distributions, enables regime identification of the measured pERK distribution. We further show qualitative assessment of the modality of the response distribution can constrain the parameter range within an operating regime. Both R_IQR_ and R_M_ being experimental observables makes the proposed method suitable for assessing signal modulation capability of an enzymatic cascade of interest.

## Introduction

Enzymatic cascades such as MAPK pathway are crucial, ubiquitously conserved, building-block in a signaling network [1,2]. Phenotypic response of a cell to a certain stimulus is known to be governed by the response of such cascades [3]. Insights into the distinct steady-state behavior of an enzymatic cascade can be obtained by placing the dose-response curve --- the relationship between the upstream kinase and downstream phosphorylated substrate concentrations --- into different operating regimes [4]. This identification of the regimes is currently amenable to only deterministic framework, that is, population-averaged behavior. However, cell-to-cell variability inherently present in a population of cells plays crucial roles in cellular decision-making process resulting in a certain phenotype [5]. Therefore, it is necessary to find the operating regimes based on protein abundance information available at the ensemble-level.

Four operating regimes of a single enzymatic cascade, viz., hyperbolic, signal-transducing, threshold-hyperbolic, ultrasensitive (saturated) represent characteristically different behaviors based on the four combinations of saturated and unsaturated level of the kinase and phosphatase [4]. These have been identified by contrasting the dose-response curves, arrived at by using total quasi-steady state approximation (tQSSA), in different regimes with the corresponding nominal profiles. Recently, using different approximations of tQSSA model, Straube [6] classified the steady-state behavior by considering relative concentrations of the upstream kinase and substrate in addition to the deciding aspect used by Gomez-Uribe et al [4]. Moreover, Lipshatz [7] showed that the steepness of the dose-response curve is controlled by the relative availability of the kinase and phosphatase. On the other hand, several experimental studies in the past suggest that the enzymatic cascade may display different steady-state behavior at ensemble-level when compared with that for an average population [8,9]. Attempts to understand such a behavior using modeling framework typically considered variation in the dose-response curve steepness [10,11], constraints on the availability of upstream kinase [7], variation in the effective stimulus strength [12].

Detection of the abundance of activated proteins, such as pERK in a MAPK enzymatic cascade, at an ensemble level is typically achieved by quantitatively capturing fluorescence emitted by an appropriate (non-plasmid) reporter [3,8,9,13]. These reporter-based detection techniques hinges on cell-fixation and permeabilization for permitting entry of protein specific antibodies. As a result, they are amenable for capturing continuous time-trajectories of ensemble of cells and are rather suitable for obtaining snapshots of the distribution of abundance at the discrete sampling points [14]. Since the enzymatic cascade is present in each of the cells, the entire population could be responding according to a certain operating regime. We therefore ask a question if the underlying operating regime that dictates the activated protein abundance in a population of cells can be directly inferred from such ensemble-level distribution data.

In order to address this question, in this study, we consider identifying the operating regimes of a single enzymatic cascade exhibited at ensemble-level. Using Michaelis-Menten approximated mathematical model of the single enzymatic cascade as a basis, we systematically assess the nature of relationship between representative signatures of the distributions of upstream kinase (pMEK) and of the downstream phosphorylated substrate (pERK). We show that this relationship can be reliably capitalized upon for identifying the operating regimes from distribution information constrained by the experimentally measured ensemble-level data of pMEK and pERK in a model Jurkat E6.1 cells stimulated with different PMA concentrations. Using this relationship, we suggest the regime in which the measured experimental distributions may lie in.

## Results

Ensemble-level behavior of pERK response is known to be different compared to the population-average behavior [8]. In order to assess the relationship between the ensemble-level behavior of an upstream kinase and the downstream activated substrate, we consider the classical single phosphorylation-dephosphorylation enzymatic cascade involving phosphorylated-MEK (pMEK) acting as a kinase for transitioning of the inactive form of ERK its active form (pERK) via phosphorylation (Fig. 1). A phosphatase P reverts the active to inactive form through a dephosphorylation reaction. Since the primary objective of the study is deciphering operating regime using ensemble-level snapshot data, we first consider experimental detection of the discrete-time fluorescence distributions as a response to Phorbol Myristate Acetate (PMA) stimulation. Note that PMA stimulation is often used as an experimental model to study ERK activation via Protein kinase C (PKC), a central event in Toll like receptors (TLR) and T cell receptor complex(TCR/CD3) pathways [15–19] .

**Figure 1:**
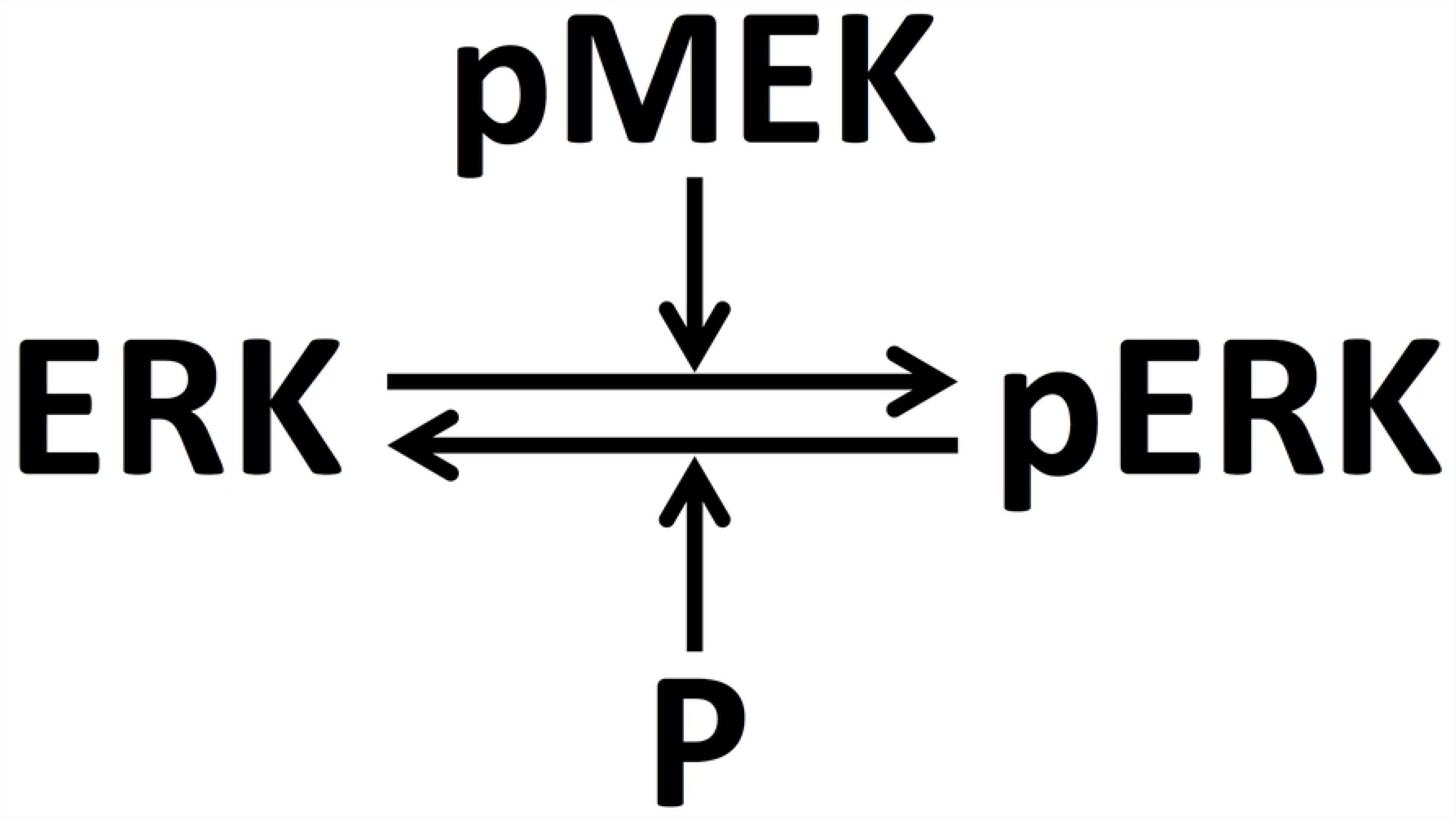
Schematic of a single enzymatic cascade.

### Discrete-time snapshot distributions of pMEK and pERK

Using Jurkat E6.1 as model cell line, we simultaneously detected the instantaneous snapshots of fluorescence levels of pMEK and pERK in a population of cells exposed to 1ng/ml (T1), 100 ng/ml (T2), 1000 ng/ml (T3) concentrations of PMA for 0 to 30 min. For this purpose, we used a dual-staining flow cytometry reporter assay (Materials and methods). Instrument and handling variability were minimized by employing a high-throughput fluorescent cell barcoding (FCB) before staining and acquisition [20] (Materials and Methods).

Ensemble-level dynamics along with the corresponding mean fluorescence intensities (MFIs) and standard deviation (SD) for both pMEK and pERK for all three treatment conditions is presented in Fig 2 – replicates in Fig. S1-I and Fig. S1-II. Note that the pMEK and pERK distributions for unstimulated cells (control) are in Figs 2A and 2D, respectively. While pMEK levels for T1 was similar to that of control (Fig. 2A), there is significant activation for treatments T2 and T3. Dynamics of the distributions, MFI and SD (discounted for deviation across replicates) suggests that cells may have reached a stationary-state within 30 mins for all three treatments. Thus, henceforth, for the rest of the results reported, unless otherwise explicitly mentioned, as steady-state, we assume the distribution obtained for at 30 mins.

**Figure 2:**
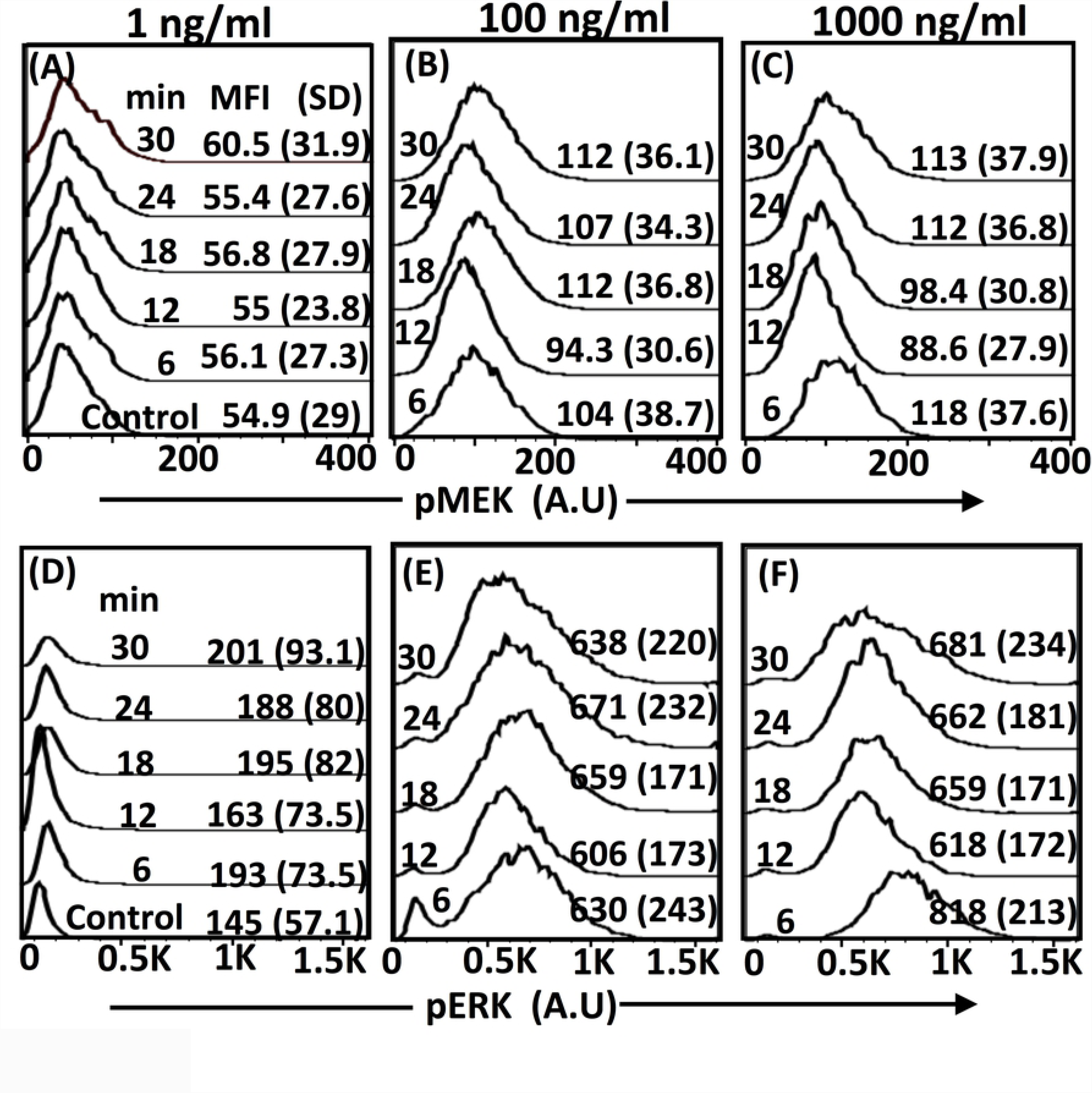
Normalized distributions of pMEK and pERK in JurkatE6.1 cells exposed to 1ng/ml (A & D), 100 ng/ml (B & E), 1000 ng/ml (C & F) PMA for different durations. The cells were dual-stained with Alexa488-pERK(1/2) and Alexa647-pMEK. Distribution corresponding to the control (unstimulated) is in the panel for 1 ng/ml (A & D). MFI and SD, respectively for every distribution corresponds to its mean fluorescence intensity and the standard deviation. Replicates are in Fig S1.

### Mathematical model: Ensemble-level behavior

We next consider a mathematical model of the single-enzymatic cascade (Fig. 1) with biochemical reactions

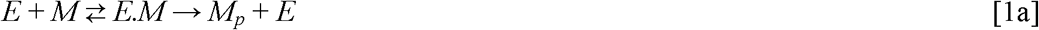

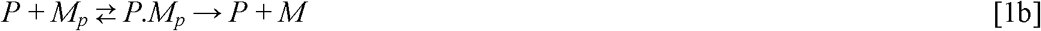

where *E, M*, and *M*_*p*_, respectively represent pMEK, ERK, and pERK. Note that *E.M* and *P.M*_*p*_ are the intermediate species for which we assume quasi-steady state. The dynamics of *M*_*p*_ is

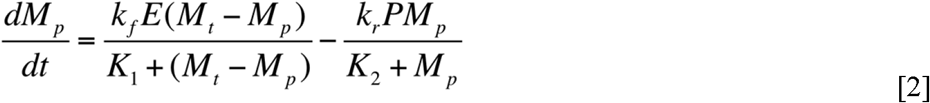

where, *k*_*f*_ and *k*_*r*_ are the forward and reverse catalytic rate constants. While *K*_*1*_ and *K*_*2*_ are Michaelis-Menten constants, *M*^*t*^ is the total substrate [10, 21]. The analytical solution of Eq. 1 capturing the steady-state behavior is

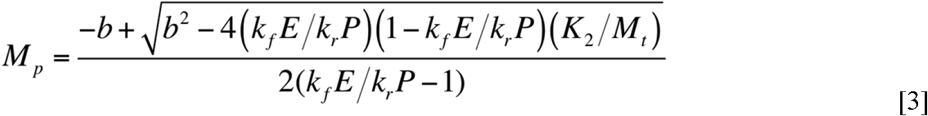

where, *b*=–(*k*_*f*_*E*/*k*_*r*_*P*–1)+(*K*_2_/*M*_*t*_)(*k*_*f*_*E*/*k*_*r*_*P*)+(*K*_1_/*M*_*t*_). For the sake of brevity, we henceforth refer to the dose-response curve dictated four operating regimes, Hyperbolic, Signal-transducing, Threshold-hyperbolic, and Saturated, respectively as H, ST, TH, and S. Note that these regimes correspond to different combinations of kinase and phosphatase being saturated or unsaturated as controlled by *K*_*1*_ and *K*_*2*_ [4].

While the regime identification at the steady state deterministic level is by comparing dose-response for a given set of parameters with a nominal standard profile ([4], Methods), such an approach, is not directly amenable for use in the snapshot experimental data such as those in Fig. 2. Moreover, for such an experimental system where snapshot data alone is available, the underlying dose-response curve is typically unavailable. Since different operating regimes have characteristically different dose-response curve, the distributions obtained at ensemble-level, which are guided by the underlying steady-state behavior, may display a qualitatively distinct behavior in different regimes. Therefore, we suppose it is rather prudent to base the identification of operating regime on both the distribution-level metric and the qualitative contrasting of the distributions of upstream enzyme E (input) and of phosphorylated substrate M_p_ (response). In the following section, we first consider the qualitative assessment of the operating regimes, and subsequently (in a future section) propose a distribution-level metric and assess its suitability.

We chose parameters for the model simulations based on those reported in Gomez-Uribe *et al*. (Table 1). We further specify in Table 1 the *K*_*1*_ and *K*_*2*_ combinations that would correspond the four different regimes. A sampling strategy (Methods) was used to generate 140000 (K_1_, K_2_) sets. Dose-response curve corresponding to each of the samples were contrasted with nominal profiles and those sets belonging to different regimes were identified (Methods). While 58.9 % of sets were placed in four different regimes, the remaining dose-response curves do not satisfy the regime-identification criteria. For the rest of this study, the fraction of parameter sets placed in their respective regimes is used for further analysis.

**Table 1:**
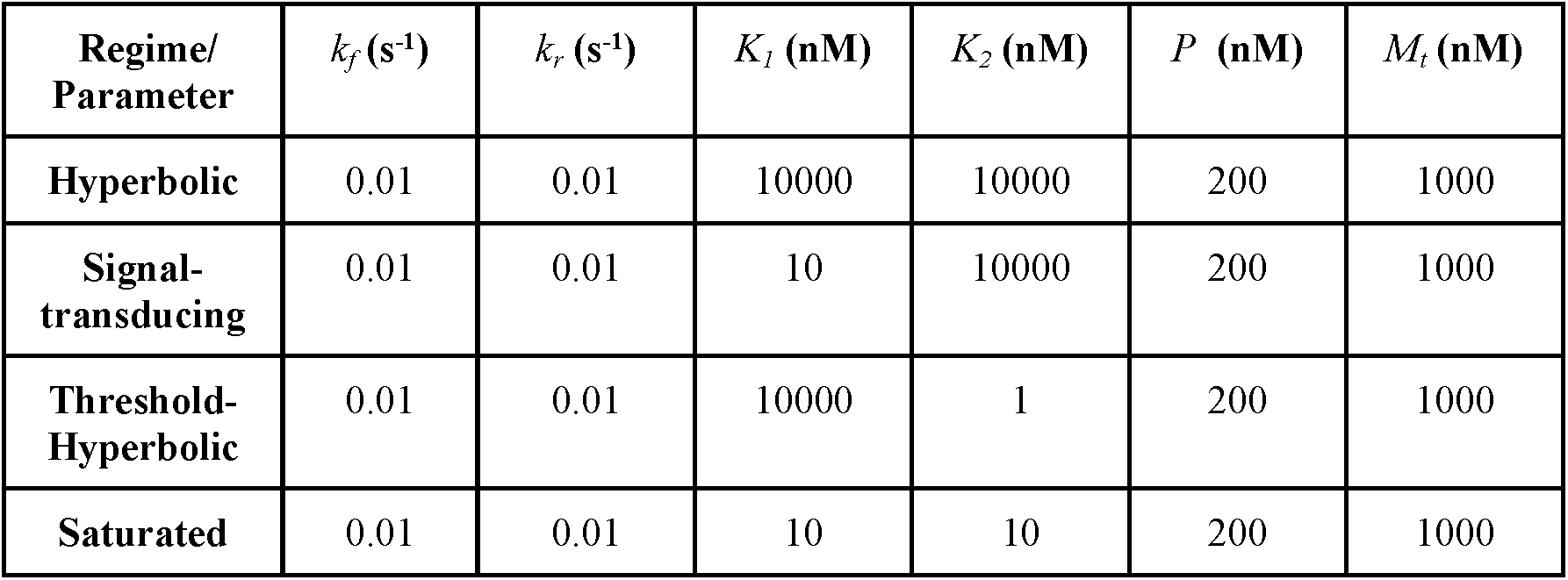
Parameters corresponding to nominal profile for the four operating regimes. These are based on those reported in Gomez-Uribe et al [4].

### Bimodal response in Signal-transducing regime

Qualitative assessment of relationship between the input and response distributions is possible only if the knowledge of all possible qualitatively different ensemble-level behaviors (within an operating regime) are known. In this section, we systematically distill out various qualitative behavior possible at ensemble-level in the four operating regimes. Variability in the response of a population of cells can be attributed to several sources such as distribution of protein levels in resting cells, variation in cell-cycle stages [22, 23]. In this study, we assume that the variability across cells in a population stems from the possibility of levels of upstream enzyme *E* being different in them, as suggested in [12]. Under these assumptions, the underlying dose-response curves in TH [13] and S [8] regimes may offer the possibility of bimodal distribution for *M*_*p*_. However, it is as yet unclear if the dose-response curves in the ST regime could permit a bimodal response behavior. In order to assess this possibility, we first consider two distinct dose-response curves in the ST regime, *viz*., for *K*_*1*_=25 nM and *K*_*1*_=95 nM for a fixed *K*_*2*_=1000 nM (Fig. 3A). (Note that all other parameters for these two are same, and also that in ST regime, kinase and phosphatase, respectively are saturated and unsaturated.)

**Figure 3:**
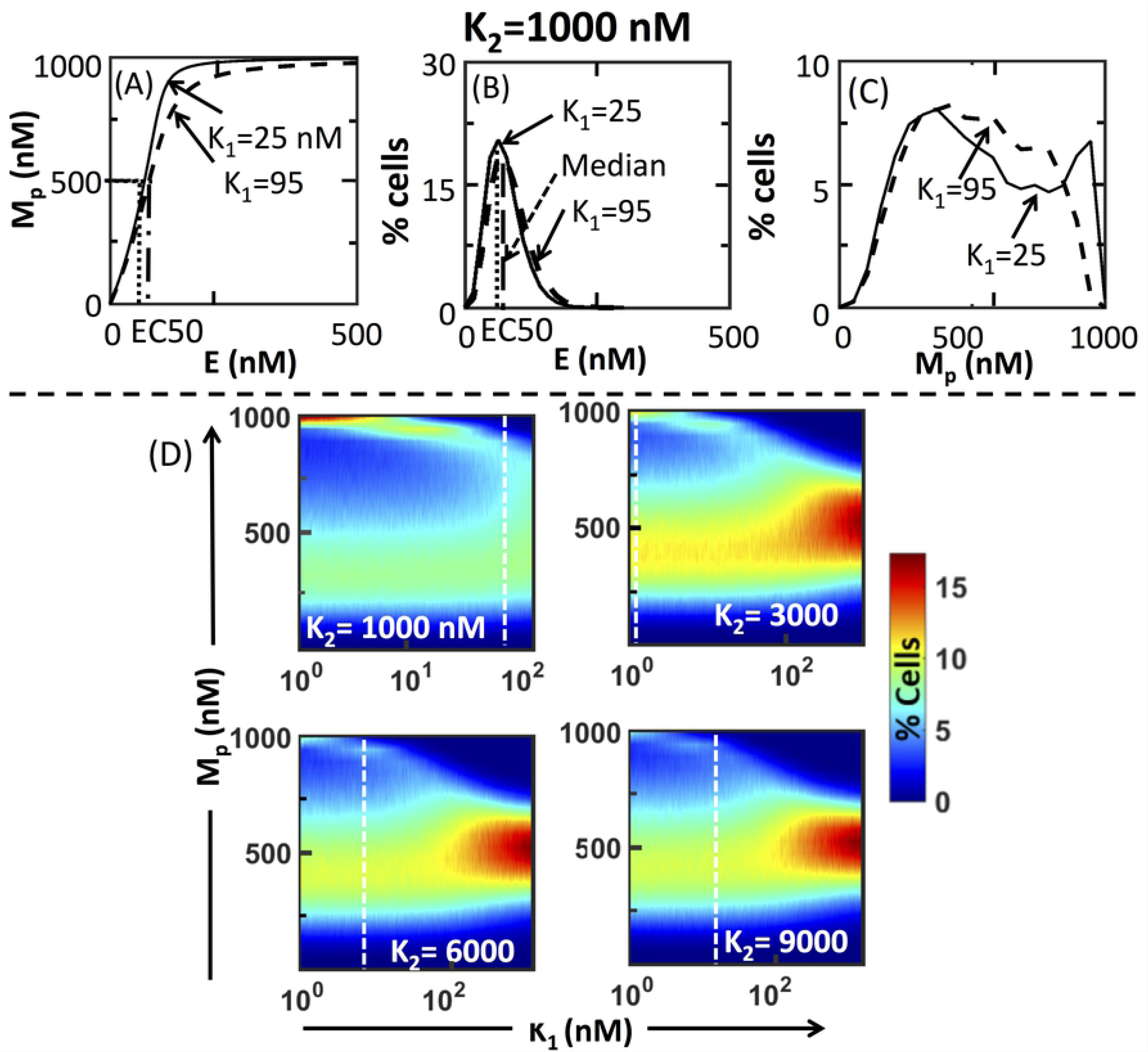
Signal-transducing regime can permit both unimodal and bimodal response. (A) Dose-response curves corresponding to (*K*_*1*_,*K*_*2*_)=(25,1000) and (95,1000), both in ST regime. (B) Input kinase distribution (with EC50 as median and shape parameter of 4.35) corresponding those profiles in (A). (C) The response distribution corresponding to the profiles in (A) subjected to the input kinase distribution in (B). (D) Normalized pERK distributions in the ST regime in planes of *K*_*1*_ and *M*_*p*_ for four different fixed *K*_*2*_. White dashed line separates the (*K*_*1*_, *M*_*p*_) plane based on unimodal or bimodal nature of the normalized response distribution (*M*_*p*_) using Hartigan’s dip test.

We next introduce ensemble-level features of the two dose response curves. The first step towards superimposing ensemble-level features on the dose-response curve is identification of input distribution. Based on the approach in Birtwistle *et al*. [13], we deduced that the experimental pMEK abundance (at 30 mins) in Fig. 2 can be approximated to gamma distribution (Text S1-I) with the estimated shape parameter in the range (4.16, 8.65). We therefore considered a gamma distribution for the upstream kinase (*E*) with a nominal value of shape parameter of 4.35. (Note that we later consider a wider range of shape parameter and demonstrate its influence on the predictions.) Further, in order to ensure a poised representation of the entire M_p_ concentration range, we placed the median of the distribution at E_1/2_, the concentration of *E* corresponding to *M*_*p*_=500 nM, for the specified parameters. (Note that the value *E*_1/2_ for a set of parameters can be directly estimated from Eq. 3. Moreover, it can be shown that the maximum *M*_*p*_ concentration permitted by Eq. 3 will be that of total *M*_*t*_ (=1000 nM) for sufficiently large concentration of *E* irrespective of the other parameters in the model.) The input distributions corresponding to the dose response curves at *K*_*1*_=25 nM and *K*_*1*_=95 nM for a fixed *K*_*2*_=1000 nM are in Fig. 3B.

We generated 20000 values for the upstream kinase from the input gamma distribution. For each of these, using the dose-response curve for *K*_*1*_=25 nM or 95 nM as a map, we found the corresponding *M*_*p*_ level. Thus, by superimposing the assumed input distributions on the dose-response curves, we generated the response (*M*_*p*_) distribution for the two cases considered (Fig. 3C). Hartigan’s dip test [24] confirmed that the distribution obtained for *K*_*1*_=95 nM is unimodal and that for *K*_*1*_=25 nM is not. This suggests the different range of *K*_*1*_ in ST regime could permit qualitatively different ensemble-level behavior.

We next assessed the presence of unimodal and bimodal M_p_ distributions in the range of *K*_*1*_ and *K*_*2*_ permitted by the ST regime. In Fig. 3D, we show the effect of *K*_*1*_ on the *M*_*p*_ distributions for different *K*_*2*_ with normalized histogram for every *K*_*1*_ captured by the heatmap with colors representing percentage of cells. Hartigan’s dip test (with p-value < 0.05) was used to separate the *K*_*1*_ parameter space into ranges that permit bimodal and unimodal distributions (dashed white line in Fig.3D). Fig. 3D clearly shows that for a wide range of K_2_, there exists two regions within the signal transducing regime that permit characteristically different ensemble-level behavior. Moreover, the range in *K*_*1*_ for which bimodal response distribution may be observed decreased with increase in *K*_*2*_. Since the response distribution could be sensitive to nature of the input gamma distribution, we considered the effect of the shape parameter on the presence of the unimodal and/or bimodal response distribution. For a wide range of shape parameters, the ST regime could permit both unimodal and bimodal response distributions (Fig S2).

We next repeated this qualitative characterisation of the response distributions (dictated by the input distributions) obtained for the other three regimes, viz., hyperbolic (Fig S3), threshold-hyperbolic (Fig S4) and saturated cases (Fig S5). While for the case of threshold-hyperbolic both unimodal and bimodal responses were observed at different *K*_*1*_ and *K*_*2*_ ranges, hyperbolic regime permits only unimodal distribution for all parameter ranges (including shape parameter of the input distribution). Note that we find only bimodal response in saturated regime. While low values of shape parameter permits only bimodal response in the threshold-hyperbolic regime, only unimodal response is exhibited for large values of shape parameter (Fig S4).

### Monotonic relationship between inter-quartile range ratio (R_IQR_) and median ratio (R_M_)

We next consider the case of the population-level approach for regime identification using the snapshot distribution. The regime identification that uses the underlying dose-response curve depends both on the levels of the upstream kinase and that of the response protein *M*_*p*_. Thus the properties of both distributions needs to be considered simultaneously in arriving at an appropriate approach.

Besides the mean and median, which reflect the average properties of a distribution, metrics that capture the variation in the protein levels across cells in a population include variance, coefficient of variation (CV) given by the ratio of standard deviation and mean. While the CV of the response distribution could be considered, the CV of a input gamma distribution is uniquely specified by the shape parameter (Text S1-I). Therefore CV cannot be used for contrasting the nature of input and response distributions. On the other hand, since gamma distribution is a skewed distribution, we choose inter-quartile range (IQR), a balanced scale estimator [25], of the input and the response distributions as representative signature for the cell-to-cell variability in the protein levels of *E* and *M*_*p*_, respectively. A ratio of the IQR of the response and the input distributions, R_IQR_ can offer insights on the extent of deviation in the variability offered by the enzymatic cascade subject to dose levels dictated by a gamma distribution.

We next assess the nature of the R_IQR_ on the median of the input distribution *E*_*m*_. Since *E*_*m*_ is equal to EC50 of dose-response curve, its value can be inferred from *K*_*1*_ and *K*_*2*_ [11]. In Fig. 4, we show, as a scatter, the dependence R_IQR_ on R_M_, the ratio of median of *M*_p_ (=500 nM) and *E*_*m*_, for each of the regimes. Note that the input distribution for a set of parameters is uniquely defined by *E*_*m*_ for a fixed shape parameter. Every point (having same color) in the scatter represents a specific *K*_*1*_ for a fixed *K*_*2*_. The relationship between the R_IQR_ and R_M_ is captured for a few distinct values of *K*_*2*_ (different colors). (Note that in Suppl Mat. Fig. S6, we present the same for a range of *K*_*2*_ with different fixed *K*_*1*_.). In the H regime (Fig 4-H, Fig S6-H), the R_IQR_ is linearly correlated to the corresponding R_M_ in the range of *K*_*2*_ (or *K*_*1*_) values. For the case of TH regime (Fig. 4-TH, Fig. S6-TH), the linear relationship between R_IQR_ and R_M_ can be observed only after discounting for small deviation. In the ST regime, while the relationship for specified value of *K*_*2*_ (Fig 4-ST) is monotonic in the given range of R_M_, the same for specified value of *K*_*1*_ (Fig S6-ST) is linear. On the contrary, the saturated regime (Fig. 4-S, Fig. S6-S) shows no correlation as such, which can be attributed to the fact that the variance of response distribution is indeterminate for saturated profiles (Detailed proof in Appendix). Moreover, the qualitative relationship between the R_IQR_ and R_M_ for H, ST and TH regimes is nearly insensitive to the variation in the shape parameter (Fig S7).

**Figure 4:**
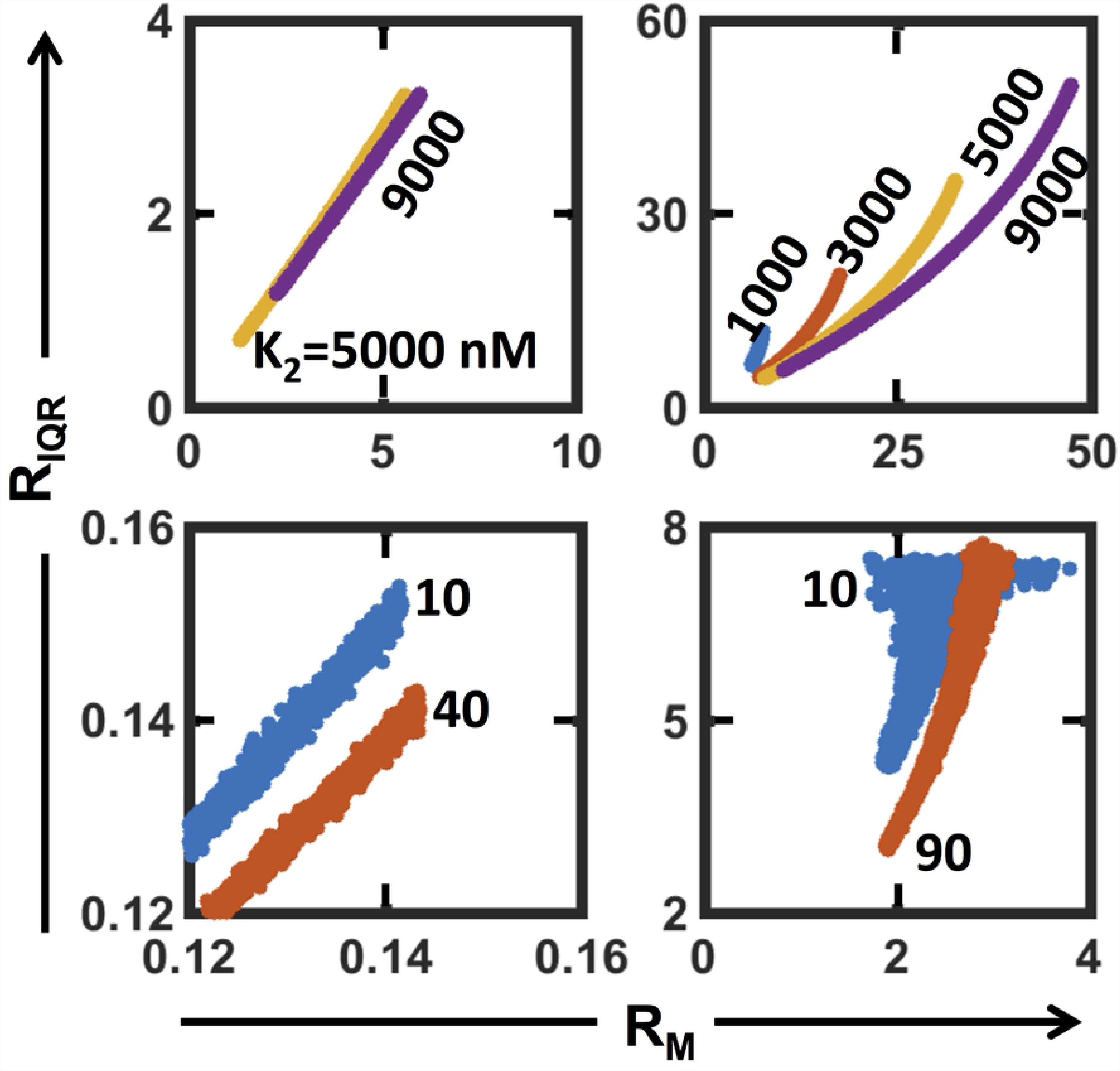
Relationship between R_IQR_ and R_M_ for the regime permitted *K*_*1*_ range at a few fixed *K*_*2*_ values for the four operating regimes. Note that similar relationship for the regime permitted *K*_*2*_ range at a few fixed *K*_*1*_ values is in Fig. S7.

Using the range permitted by the experimental data (Fig. 2) and reported in literature [13], we consider three cases of low, medium and high values for the shape parameter for input gamma distribution. In Table 2, we report the range for the R_IQR_ obtained for the four operating regimes for each of these three cases. Upon discounting for the marginal overlap in the distributions (as shown in Fig. S8), the R_IQR_ (of a given input-output distribution pair) itself can serve as a reasonable marker to identify the regime in which the underlying enzymatic cascade may lie. Note that for high shape parameter, that is, low CV of the gamma distribution, R_IQR_ range obtained in the S regime overlaps significantly with that in the ST regime. This could be due to the fact that the underlying dose-response curves in the S regime having R_IQR_ in (∼5, ∼17.26) are perhaps similar to those obtained in the ST regime but R_IQR_ being in the same range. This suggests that if the input distribution has a shape parameter around the medium and high values reported in Table 3, then R_IQR_ alone may be used to directly identify the operating regime of the underlying cascade that governs the experimental input-output distributions.

**Table 2:** Effect of shape parameter of the input gamma distribution on the lower and upper limits of R_IQR_ for the four regimes.

| Regime→<br>/Shape<br>parameter↓ | Threshold<br>Hyperbolic<br>$R_{IQR}$ | Hyperbolic<br>$R_{IQR}$ | Saturated<br>$R_{IQR}$ | Signal-Transducing<br>$R_{IQR}$ |
| --- | --- | --- | --- | --- |
| **1.56** | (0.14,0.17) | (0.83,3.24) | (2.55,6.02) | (5.85,54) |
| **4.35** | (0.12,0.15) | (0.73,2.90) | (2.6,8.03) | (5.20,54.43) |
| **11.11** | (0.11,0.13) | (0.71,3.01) | (2.66,17.26) | (5.01,54.58) |

**Table 3:** R_IQR_, R_M_ and p-Value for Hartigan’s dip test along with likely operating regime that corresponds to the input and response experimental distributions for the three PMA treatment conditions.

| **Sample** | **p-value of Hartigan's dip test** | **Median ratio ($R_M$)** | **IQR ratio ($R_{IQR}$)** | **Likely Regime** |
| --- | --- | --- | --- | --- |
| **Control** | $0.95 \pm 0.005$ | $1.60 \pm 0.03$ | $3.08 \pm 0.15$ | Hyperbolic |
| **1ng/ml PMA** | $0.99 \pm 0.003$ | $1.93 \pm 0.08$ | $3.36 \pm 0.007$ | Hyperbolic |
| **100ng/ml PMA** | $1 \pm 0$ | $6.75 \pm 0.12$ | $6.78 \pm 1.21$ | Signal-transducing |
| **1000ng/ml PMA** | $0.99 \pm 0.001$ | $6.98 \pm 0.31$ | $5.43 \pm 1.27$ | Signal-transducing |

Note that R_IQR_ metric does not distinguish the unimodal and bimodal distributions, which needs to be identified using the qualitative assessment outlined in the previous section. Thus, a combination of the qualitative assessment of the nature of the distribution -- unimodal or bimodal -- along with the quantitative metric of R_IQR_ is necessary for placing the state of the cells reflected by the input and response distributions in an appropriate operating regime. Note that placement of the distributions in the saturated regime, wherein the histograms are expected to be bimodal, can be achieved simply by elimination as the monotonic relationship between R_IQR_ and R_M_ cannot be guaranteed. R_IQR_ is purely based on the properties of response and input distributions, and does not require the availability of dose-response curve for identifying the regime that could correspond the distributions.

### Effect of PMA stimulus on the operating regime

We next implement the qualitative assessment and the relationship between R_IQR_ and R_M_ to identify the operating regime that may correspond the pMEK-pERK input-response distributions in Fig. 2. Response distributions of all three treatment conditions at 30 mins exhibit a unimodal behavior, as reflected by the Hartigan’s dip test with confidence-level greater than 95% (Table 3).

We next extracted the R_M_ and the R_IQR_ for the control and the three treatment conditions, T1, T2, and T3 (Table 2). We next assessed if the obtained (R_M_, R_IQR_) set from pMEK-pERK distribution pair for each of the treatments were in the model predicted (R_M_, R_IQR_) range that permitted monotonic behavior for H, TH, and ST. (R_M_, R_IQR_) of (1.60,3.08) and (1.93,3.36), respectively for the case of resting cells (control) and for cells exposed to 1 ng/ml (T1) suggested that the underlying cascade is operating in the hyperbolic regime. On the other hand, for other two treatment conditions of 100 ng/ml (T2) and 1000 ng/ml (T3), the corresponding (R_M_, R_IQR_) showed that the underlying cascade is likely operating in signal-transducing (ST) regime.

### Conclusion and discussion

Population-level measurements leading to a distribution of abundance of intra-cellular proteins in an ensemble of cells needed for characterising cell-to-cell variability in non-plasmid reporter based assays is available only at discrete time points [14]. Often the underlying dose-response curve, which captures the steady-state relationship between the trigger kinase and response active phosphorylated protein in the cascade, is unavailable. Operating regime, a crucial signature of the interplay between the kinase and the substrate [4, 6], currently identified using the dose-response curve cannot be deciphered when only snapshot distribution data is available. In this study, we present a novel systematic method by which such population-level snapshot experimental data can be directly used to decipher the underlying operating regime in which the cells may be in. This method systematically identifies the operating regime by juxtaposing (a) the model-based relationship between the R_IQR_, ratio of the inter-quartile range of the experimental input and response distributions and R_M_, the ratio of the medians of the two distributions, and (b) qualitative assessment of the unimodal vs bimodal nature of the response distribution. Further, the qualitative assessment suggests the possible range of parameters (Michealis-Menten constants) within the identified regime that may correspond to the biochemical reactions occuring in the cells considered. We implement the proposed method to suggest the possible operating regime corresponding to the pERK (response) and pMEK (input) distributions in Jurkat-T cells stimulated by different concentrations of PMA.

Systematic analysis of model-based response distributions governed by dose-response curve and experimental data guided upstream kinase gamma distribution revealed that signal transducing (ST) and threshold-hyperbolic (TH) regimes may permit both unimodal and bimodal response (Figs. 3, S3 and S4). The nature of the thus obtained response was catalogued for a wide range - four orders of magnitude - of two Michaelis-Menten constants (Figs 4 and S6). While ability of the cascade to permit bimodal distribution in the TH regime has been reported in literature [13], to the best of our knowledge, this is the *first* study to show that fitting superimposition of the kinase distribution on the dose-response curve can lead to bimodal behavior even in the ST regime.

The ability to decipher the operating regime from the experimental distribution data strongly hinges on the fact that the R_IQR_ exhibits monotonic relationship with the R_M_ in H, TH and ST regimes. This could be due to the fact that the unique mapping between the upstream kinase and downstream phosphorylated substrate levels is enforced by the underlying dose-response. However, no such relationship could be established in the saturated regime because the variance in this regime is likely to be indeterminate (Appendix I). R_IQR_, after discounting for overlaps, being well-separated (Table 2) for medium and high shape parameters of the input distribution corroborates the potential use of the proposed metric and method (R_IQR_ vs R_M_) for identifying the operating regime from the experimental snapshot distribution data. It is yet unclear what the relationship between R_IQR_ and R_M_ in the saturated regime is and why it is not monotonic. While the total substrate variability is not considered in this study, it can have an influence on the monotonic relationship observed. A natural extension of this study would be to systematically assess the sensitivity of the monotonic relationship on the other parameters, particularly that of total substrate concentration.

Implementation of the proposed method revealed that, at steady-state, while Jurkat E6.1 cells under resting and 1 ng/ml of PMA are likely in hyperbolic operating regime, those stimulated for 100 and 1000 ng/ml PMA exhibit response distributions that may be governed by signal-transducing behavior. Such a dose-dependent variation in the operating regime could be due to the elevated signaling in the cells caused by higher dosage leading to increased levels of the upstream kinase.

## Methods

### Cell-culture and reagents

Jurkat E6-1 cells were procured from National Centre for Cell Science(NCCS),Pune(India), and cultured in RPMI1640 supplemented with Fetal bovine serum (10%v/v),2mM/L L-glutamine and 1%antibiotic-antimycotic solution,(procured from HiMedia, India) and were maintained at 37°C in humidified 5% CO2. Cells were serum starved overnight prior to harvesting for further use. Phorbol Myristate Acetate(PMA) and Alexa 350, respectively were procured from Sigma Aldrich and Invitrogen/ThermoS. Primary conjugated Alexa488-ERK1/2(pT202/pY204) and Alexa647-MEK1/2(pS218) antibodies were procured from BD BioSciences.

### Flow cytometry based single-cell level fluorescence detection

One million harvested cells were stimulated with PMA for the indicated concentrations(1ng/ml, 100ng/ml and 1000ng/ml) for 30 min, fixed using 4% Paraformaldehyde(PFA) for 10 min at 37°C. Cells were then washed and permeabilized using 80% methanol for at least 15min on ice. Stimulated cells were stained and multiplexed using fluorescent cell barcoding (FCB) [26] before acquisition on BD FACS Aria. Captured data was first corrected for morphology variation [27], then deconvoluted and analysed using FlowJo [20].

### Parameter sampling

Based on the nominal values of (*K*_*1*_, *K*_*2*_) in Table 1 and uniform distribution, 140000 sets of (*K*_*1*_, *K*_*2*_) were generated using stratified random sampling to represent adequately all four combinations of saturated and unsaturated states of the kinase and phosphatase. Details are in Text S1-II.

### Regime identification

For every (*K*_*1*_, *K*_*2*_), and all other parameters fixed, the dose-response curve was generated using the model steady-state mapping (Eq. 3) between the upstream kinase and the phosphorylated substrate. Nominal dose-response curve (profile) was constructed for each regime using the parameters in Table 1. Relative euclidean distance (*d*_*c*_) between the dose-response curve for a certain (*K*_*1*_, *K*_*2*_) and the nominal profile of the four regimes was estimated. Details are in Text S1-III. For a particular (*K*_*1*_, *K*_*2*_), the regime for which *d*_*c*_ < 10%, the dose-response curve corresponding to that parameter combination was earmarked for that regime.

## Acknowledgements

We thank Department of Science and Technology, Government of India for funding this study. FACS Central facility, IIT Bombay is acknowledged for access. One of the authors (AP) is partially funded by UGC, Government of India.

## Appendix I Response distribution variance in saturated regime

Based on the stationary distribution of the phosphorylated substrate for a single-enzymatic cascade (Levine, Kueh, and Mirny 2007), in the saturated regime, the mean and variance of the distribution, respectively are

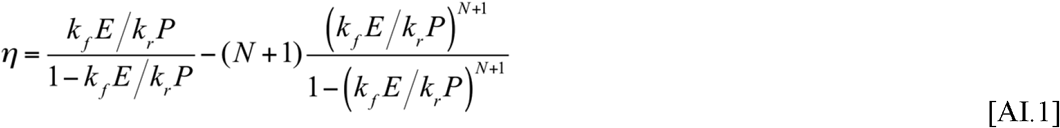

and

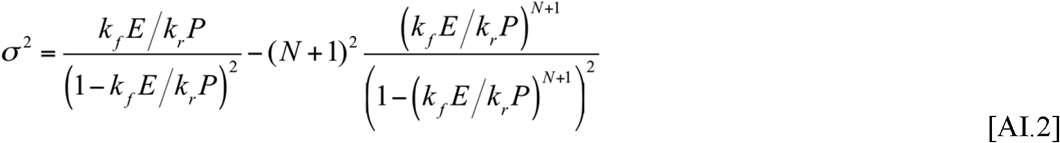

where, N is the ensemble size. All other symbols are as in Eq. 1. Using E1/2 for the (K1,K2) samples in saturated regime, the range for *k*_*f*_ /*k*_*r*_*P* is [0.9,1]. Thus, for this range, the second term in Eqs AI.1 and AI.2 will vanish for the ensemble size of 20000 considered in this study. As a result, the mean and variance, respectively are

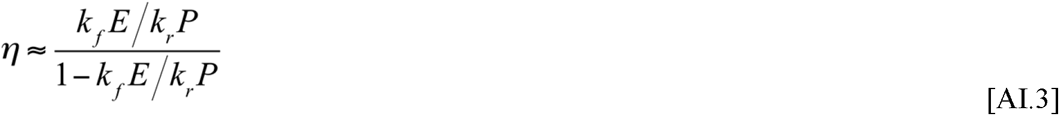

and

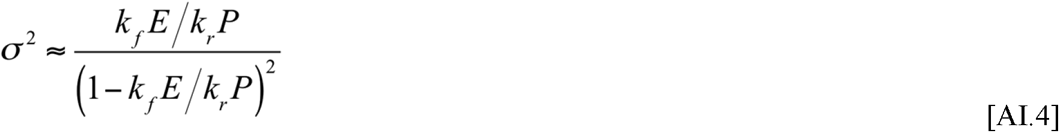

For the case nominal profile of saturated regime, wherein both phosphorylation and dephosphorylation reactions are operating under saturation conditions, *k*_*f*_*E*/*k*_*r*_*P* ≈ 1. Thus, for the dose-response curves approaching nominal profiles in saturated regime, standard deviation of the distribution is likely to be indeterminate.

## Figure captions

*Figure S1*: Two replicates (panels I and II) of the normalized distributions of pMEK and pERK in Jurkat E6.1 cells reported in Fig. 2 of main text.

*Figure S2*: Effect of shape parameter *a* and *K*_*1*_ on pERK distributions on *K*_*1*_ in ST regime for a fixed *K*_*2*_. For the sake of comparison, distributions from Fig. 3D (for a=4.35) are repeated here.

*Figure S3*: Effect of shape parameter *a* and *K*_*1*_ on pERK distributions on *K*_*1*_ in H regime for a fixed *K*_*2*_.

*Figure S4*: Effect of shape parameter *a* and *K*_*1*_ on pERK distributions on *K*_*1*_ in TH regime for a fixed *K*_*2*_.

*Figure S5*: Effect of shape parameter *a* and *K*_*1*_ on pERK distributions on *K*_*1*_ in TH regime for a fixed *K*_*2*_.

*Figure S6*: Relationship between R_IQR_ and R_M_ for the regime permitted *K*_*2*_ range at a few fixed *K*_*1*_ values for the four operating regimes. Note that similar relationship for the regime permitted *K*_*1*_ range at a few fixed *K*_*2*_ values is in Fig. 4.

Figure S7: Effect of shape parameter on the relationship between R_IQR_ and R_M_. Note that in the ordinate captures the effective slope of the monotonicity between R_IQR_ and R_M_, as estimated using a linear fit.

*Figure S8*:-Distribution of R_IQR_ (in the form of boxplots) for the four regimes for low, medium and high values of shape parameter *a*.

